# Six species of non-tuberculosis mycobacteria carry non-identical 16S rRNA gene copies

**DOI:** 10.1101/302836

**Authors:** Keita Takeda, Kinuyo Chikamatsu, Yuriko Igarashi, Yuta Morishige, Yoshiro Murase, Akio Aono, Hiroyuki Yamada, Akiko Takaki, Satoshi Mitarai

## Abstract

Non-tuberculosis mycobacteria (NTM) can carry two or more 16S rRNA gene copies that are, in some instances, non-identical. In this study, we used a combined cloning and sequencing approach to analyze the 16S rRNA gene sequences of six NTM species, *Mycobacterium cosmeticum, M. pallens, M. hodleri, M. crocinum, M. flavescens*, and *M. xenopi*. The approach facilitated the identification of two distinct gene copies in each species. The two *M. cosmeticum* genes had a single nucleotide difference, whereas two nucleotide polymorphisms were identified in *M. hodleri, M. flavescens*, and *M. xenopi. M. pallens* had a difference in four nucleotides and *M. crocinum* in 23. Hence, we showed that the six NTM species possess at least two non-identical 16S rRNA gene copies.

**Importance:** The presence of multiple 16S rRNA gene copies with nucleotide polymorphisms represents a challenge for species identification using 16S rRNA as the target sequence. Our analysis was focused on six NTM species, *M. cosmeticum, M. pallens, M. hodleri, M. crocinum, M. flavescens*, and *M. xenopi*. As a result, we generated the full-length sequences of two non-identical 16S rRNA copies for each NTM species. The data will be helpful for the sequence analysis of specimens or other samples.

## Introduction

Non-tuberculosis mycobacteria (NTM) comprise over 190 species of the genus *Mycobacterium* on the “List of prokaryotic names with standing in nomenclature” (http://www.bacterio.net/mycobacterium.html; accessed on April 13, 2018). NTM species are classified based on sequence comparisons of select housekeeping genes, such as 16S rRNA, *rpoB*, and *hsp65*, and the 16S-23S rRNA internal transcribed spacer region (1–10). Several commercial tests are available that use these genomic sequences for species identification (11–15). However, some NTM species cannot be identified using these tests because of insufficient genome sequence information.

Specifically, Chikamatsu et al. reported failed attempts for the identification of several NTM species using the PyroMark Q24 test kit (QIAGEN, Tokyo, Japan), which is based on pyrosequencing (16). Ambiguous bases were found within the 16S rRNA gene in six NTM species, *Mycobacterium cosmeticum, M. pallens, M. hodleri, M. crocinum, M. flavescens*, and *M. xenopi* (16). Direct sequencing using the Sanger method suggested that these species carry two non-identical 16S rRNA gene copies. In general, rapid-growing mycobacteria carry two copies of the 16S rRNA gene (17), whereas, with a few exceptions, slow-growing mycobacteria possess only one copy (18). For example, it was reported earlier that isolates of the slow-growing *M. terrae* complex (19) and *M. celatum* (20) harbor two non-identical copies of the 16S rRNA gene. Here, we applied a combined cloning and sequencing approach to unequivocally determine the copy numbers and complete sequences of all 16S rRNA genes of the six NTM species investigated earlier (16), *M. cosmeticum, M. pallens, M. hodleri, M. crocinum, M. flavescens*, and *M. xenopi*.

## Materials and Methods

### Bacterial strains

*M. cosmeticum* JCM14739, *M. pallens* JCM16370, *M. hodleri* JCM12141, and *M. crocinum* JCM16369 were obtained from the Japan Collection of Microorganisms (JCM, Ibaraki, Japan). *M. flavescens* ATCC14474 and *M. xenopi* ATCC19250 were acquired from the American Type Culture Collection (ATCC, Manassas, VA). All strains were initially grown on 7H10 agar and then cloned from single colonies. The isolates were sub-cultured in 2% Ogawa medium at 37 °C.

### DNA extraction

Bacterial DNA was extracted using the Isoplant Kit (Nippon Gene Co., Ltd, Toyama, Japan). Briefly, one inoculation loop (approximately 10 μl) of fresh colonies grown on Ogawa medium were suspended in 300 μl of extraction buffer and again suspended in 150 μl of lysis buffer for a 15-min incubation at 50 °C. Genomic DNA was extracted with sodium acetate (pH 5.2) on ice for 15 min. After centrifugation (12,000 ×g, 15 min at 4 °C), the upper phase was transferred to a new tube and the genomic DNA was precipitated with 70% ethanol. The DNA pellet was dissolved in 50 μL TE buffer (10 mM Tris-HCl, 1 mM EDTA).

### Cloning

The target 16S rRNA genes from each bacterial DNA preparation were amplified with primers 285 (5′ GAG AGT TTG ATC CTG GCT CAG 3′) and rp2 (5′ ACG GCT ACC TTG TTA CGA CTT 3′) yielding the almost complete 16S rRNA gene (1, 21). In brief, 25 μl of a mixture containing ExTaq HS (TaKaRa Bio Inc., Shiga, Japan), 2.5 mM dNTP mixture, 10 μM of each primer, and 5 μL template DNA was used for PCR. Amplification was performed in a GeneAmp PCR System 9700 (Applied Biosystems, Foster City, CA) using 30 cycles of 30 s at 94 °C, 30 s at 60 °C, and 90 s at 72 °C. Then, the PCR products were purified and cloned using a TOPO TA Cloning Kit (Invitrogen, USA). In brief, the PCR products, salt solution, water, and TOPO® vector using vaccinia topoisomerase I were mixed at room temperature (22–23 °C) and incubated for 30 min. The recombinant TA cloning mixes were incubated with *E. coli* competent cells (DH5 alpha) for 30 min on ice to perform the transformation. The process was stopped by incubating the samples at 42 °C for 60 s (heat shock), immediately followed by an incubation on ice. Super optimal broth with catabolite repression (SOC) was added to the samples and incubated at 37 °C for one hour.

The competent cells were cultured on Luria-Bertani (LB) agar supplemented with 2 mg of X-gal. Ten white colonies of each transformation were picked from the LB agar and individually cultured in LB broth. Plasmid DNA was isolated and purified using the minipreparation Flexiprep Kit (Amersham Biosciences, Buckinghamshire, England) and a column method with the FastGene Gel/PCR Extraction Kit (Nippon Genetics Co., Ltd, Tokyo, Japan).

### Sequence analysis

The sequencing of each 16S rRNA clone was performed using the primers M13 Forward (5′ GTA AAA CGA CGG CCA GT 3′), M13 Reverse (5′ CAG GAA ACA GCT ATG AC 3′) and 264 (5′ TGC ACA CAG GCC ACA AGG GA 3′) with the BigDye Terminator Cycle sequencing kit ver. 3.1 (Applied Biosystems) in an ABI 3500 Genetic Analyzer (Applied Biosystems). Finally, the sequences (approximately 1,500 bp each) of the 10 clones of each species (approximately 1,500 bp each) were aligned and further analyzed using the software package Molecular Evolutionary Genetics Analysis Ver. 7 (22).

## Results

The sequence alignments led to the identification of 2 non-identical 16S rRNA copies for each of the six NTM species. The results have been deposited with GenBank under accession number as follow: *M. cosmeticum* (MH169224 and MH169226), *M. pallens* (MH169208 and MH169209), *M. hodleri* (MH169216 and MH169217), *M. crocinum* (MH169218 and MH169219), *M. flavescens* (MH169220 and MH169222), and *M. xenopi* (MH169221 and MH169241). The nucleotide polymorphisms are shown in Figure 1. *M. cosmeticum* had a single nucleotide difference between the two sequences. Two-nucleotide differences were found in *M. hodleri, M. flavescens*, and *M. xenopi. M. pallens* had a difference in four nucleotides and *M. crocinum* in 23.

**Figure I.**
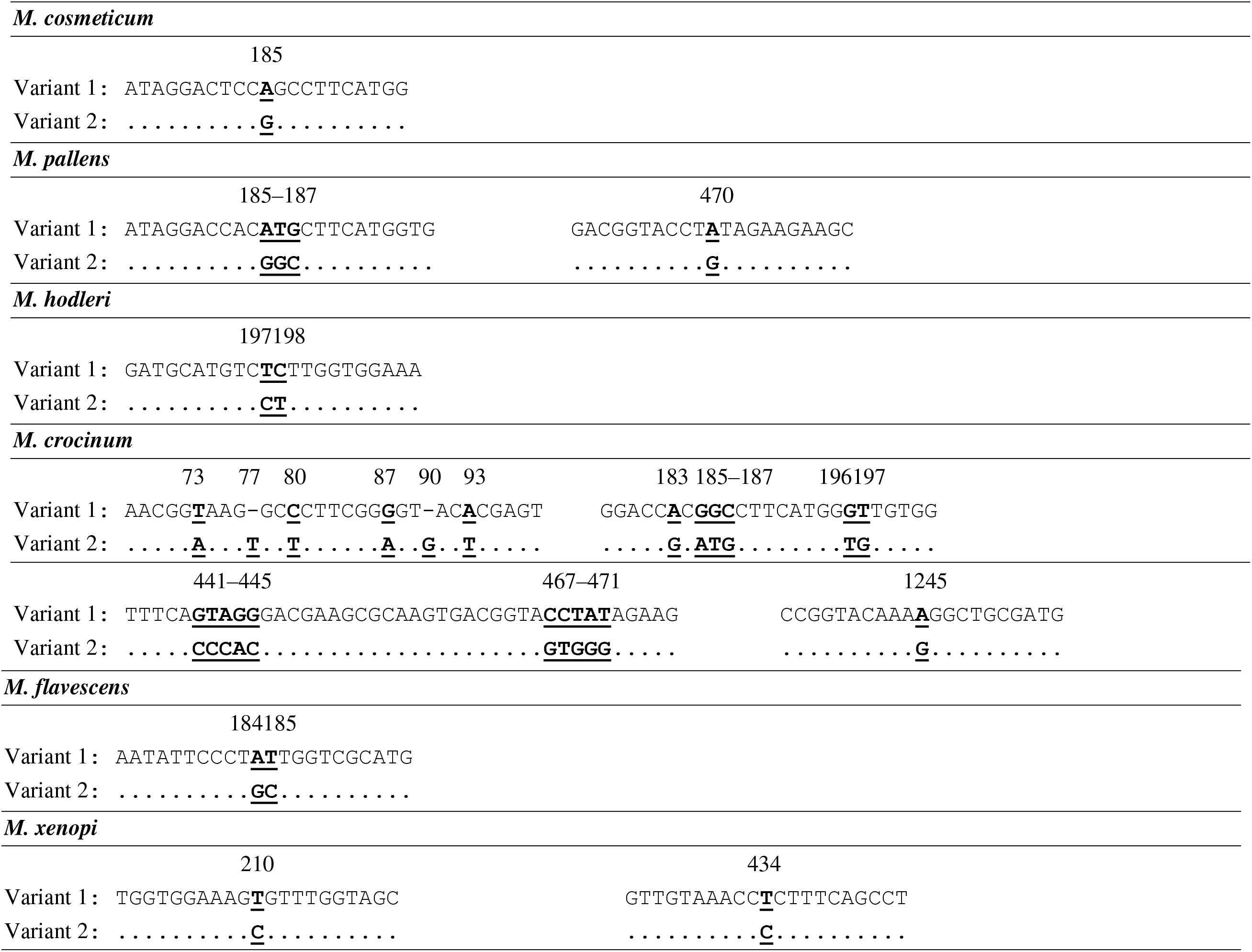
Nucleotide polymorphism between two non-identical 16S rRNA sequences of six NTM species. Nucleotide positions were derived from an alignment with the 16S rRNA gene of *M.tuberculosis* H37Rv ATCC272 (GenBank accession: NC_000962) M. cosmeticum

## Discussion

It is well documented that some species of the genus *Mycobacterium* harbor multiple 16S rRNA gene copies with distinct sequences (16, 19, 20, 23–26). In this study, the cloning experiments targeting 16S rRNA genes facilitated the identification two distinct copies in six NTM species, *M. cosmeticum, M. pallens, M. hodleri, M. crocinum, M. flavescens*, and *M. xenopi*. Each strain was re-isolated from a single colony, and each of the two 16S rRNA gene copies was reproducibly obtained from multiple clones derived from these isolates. In addition, the nucleotide polymorphisms observed for the species-specific gene copies were supported by earlier findings obtained by direct sequencing, e.g., the nucleotide position 185 of the 16S rRNA copies in *M. cosmeticum* was A or G by our cloning-sequencing experiment (Figure 1), but a mixture of A and G at this position had been indicated earlier by direct Sanger sequencing (16). Our new data were confirmed by multiple clones per strain to minimize the possible impact of technical sequencing errors.

Chilia et al. reported earlier that sequences obtained from clones may be more definitive than sequence data obtained from direct sequencing (23). Indeed, our sequence data obtained from the cloning experiment unequivocally established the existence of two non-identical gene copies per species whereas the earlier direct sequencing only suggested a polymorphism based on sequence ambiguities. Hence, it is recommended to avoid direct sequencing for species identification if there are non-identical genomic copies of the target sequence.

However, our study could not reveal whether the NTM species had more than two 16S rRNA gene copies per genome. It is possible that one genome carries several 16S rRNA copies with identical sequences. This problem might be resolved by whole-genome sequencing (WGS). A database search revealed that *M. cosmeticum* DSM 44829 has two 16S rRNA genes (GenBank accession: NZ_CCBB010000003.1), *M. flavescens* strain M6 has three (GenBank accession: NZ_MIHA00000000.1), and *M. xenopi* has one (Strain DSM 43995, GenBank accession: LQQB01000023.1) or two (Strain RIVM700367, GenBank accession: NZ_AJFI01000116.1). These WGS data were obtained by shotgun sequencing, which also has limitations regarding the identification of identical or almost identical gene copies (27–29). Hence, cloning along with sequencing is still required, but an improvement in WGS data accuracy will be obtained in near future by implementing long-read sequencing using the single molecular, real-time sequencing technology (28, 29). Currently, the exact 16S rRNA gene copy number is not yet known for *M. pallens, M. hodleri*, and *M. crocinum*, requiring further analysis.

In this study, we established the existence of two 16S rRNA gene copies for each of the six NTM species examined. However, the detected nucleotide polymorphism can be a challenge for species identification using 16S rRNA sequencing. The identification of the two non-identical 16S rRNA copies in NTM species will be helpful for sequence analyses of specimens or other samples and sequencing efforts.

